# NCK adaptor proteins regulate clathrin-coated pit dynamics and EGF-stimulated PI3K-Akt signaling

**DOI:** 10.1101/2025.10.24.684344

**Authors:** Karolina Zak, Nupur Dave, Ayshin Mehrabi, Michael G. Sugiyama, Costin N. Antonescu

## Abstract

The epidermal growth factor (EGF) receptor (EGFR) promotes cell growth, proliferation, and survival. EGF binding to EGFR activates receptor kinase activity, leading to membrane recruitment of class IA phosphoinositide-3-kinase (PI3K), thus activating Akt signaling. Clathrin-coated pits (CCPs) are plasma membrane endocytic structures that are enriched in signaling intermediates and regulate in EGF-stimulated Akt activation. The mechanisms and impact of recruitment of certain signaling molecules within the PI3K-Akt pathway to CCPs remains poorly understood. Using total internal reflection fluorescence microscopy (TIRFM), we observed that the adaptor proteins Nck1 and Nck2 are enriched within CCPs and regulate early CCP formation and maturation. Nck1 and Nck2 each support EGF-stimulated Akt phosphorylation. Notably, EGF stimulation triggers Nck-dependent enrichment of PI3K within CCPs, and perturbation of Nck adaptors suppressed cell proliferation and survival. This study identifies novel functions for Nck adaptor proteins in EGFR-mediated recruitment of PI3K-Akt signals within CCPs, Akt activation, and cell physiology.

## Introduction

The epidermal growth factor receptor (EGFR) is a receptor tyrosine kinase (RTK) that promotes cell growth, proliferation, and survival (Lemmon and Schlessinger, 2010; Wee and Wang, 2017; Orofiamma *et al*., 2022). In addition to its impactful roles in development and tissue homeostasis, aberrant activation of EGFR drives progression of several types of cancer (Sigismund *et al*., 2018; Levantini *et al*., 2022). For example, the level of EGFR present on the surface of tumor cells can heighten their responsiveness to growth factors originating from nearby healthy cells or the tumor itself (Normanno *et al*., 2006).

Binding of ligands such as the epidermal growth factor (EGF) to EGFR triggers activation of the intrinsic kinase activity of the receptor, in turn eliciting phosphorylation of tyrosine residues on the receptor’s c-terminus. These phosphorylated tyrosines are part of motifs that serve as binding sites for SH2 or PTB domains found on adaptor proteins which assist in transmitting a cascade of signals downstream of this receptor (Mitsudomi and Yatabe, 2010). In turn, signal transducers bind to these adaptors, initiating multiple signalling pathways that ultimately lead to the activation of Akt, Erk, and other pathways.

Activation of Akt upon EGF-stimulation requires phosphorylation of Y1068 on EGFR, part of a motif which binds to the SH2 domain of Grb2. This ensures recruitment of Grb2-associated binder 1 (Gab1), which associates with Grb2 via an SH3-proline rich domain (PRD) interaction. The subsequent Gab1 phosphorylation leads to binding and activation of the regulatory subunit of Class 1A phosphatidylinositol-3-kinase (PI3K) and production of phosphatidylinositol-3,4,5-triphosphate (PIP_3_) from phosphatidylinositol-4,5-biphosphate (PI(4,5)P_2_) (Mattoon *et al*., 2004; Sugiyama *et al*., 2019). PIP_3_ then promotes membrane binding and activation of Akt, with full activation of Akt achieved by phosphorylation on T308 and S473 by PDK1 and mTORC2, respectively.

There are three isoforms of Akt, Akt2, and Akt3, that are differently activated by phosphoinositides. While Akt1 and Akt3 are directly recruited and activated by PIP_3_, Akt2 is recruited by phosphatidylinositol-3,4-bisphosphate (PI(3,4)P_2_) that is produced from PIP_3_ by the phosphatase SHIP2 (Goulden *et al*., 2018; Liu *et al*., 2018). From this emerges a model in which the activation of Akt signaling is regulated through various pivotal molecular checkpoints, from ligand binding of EGFR to Gab1, Grb2, PI3K, and SHIP2, each which may contribute to Akt-isoform specific regulation.

Work from our group and others showed that full Akt activation also requires plasma membrane clathrin-coated pits (CCPs), structures well-known to also mediate endocytosis (McMahon and Boucrot, 2011; Kaksonen and Roux, 2018; Mettlen *et al*., 2018). Clathrin, adaptors, and many other proteins assemble on the inner leaflet of the plasma membrane to trigger formation CCPs, which allows coupling of recruitment of cargo proteins such as EGFR to membrane budding. Following a maturation period, the eventual scission of CCPs leads to production of intracellular vesicles. We uncovered that perturbations of clathrin that prevent CCP formation, lead to suppression of EGF-stimulated Akt phosphorylation (Garay *et al*., 2015). In contrast, perturbations of dynamin, which allow CCP formation and receptor recruitment but blunts vesicle formation, did not impact EGF-stimulated Akt phosphorylation. Other studies also found a key role for clathrin in regulation of EGF-stimulated Akt phosphorylation (Sigismund *et al*., 2008; Leyton-Puig *et al*., 2017; Rosselli-Murai *et al*., 2018; Pascolutti *et al*., 2019; Alfonzo-Méndez *et al*., 2022).

In probing the mechanism by which clathrin structures may regulate EGFR signaling, we recently showed that CCPs serve to recruit TOM1L1 and Fyn, leading to enhancement of the localization of SHIP2 to CCPs and selective regulation of Akt2 phosphorylation (Cabral-Dias *et al*., 2022). This work suggests that PIP_3_ produced by PI3K is modified by clathrin-localized SHIP2 to elicit clathrin-dependent activation of Akt2. We previously showed that Gab1 was also enriched within CCPs (Garay *et al*., 2015), while others reported that the signal-terminating lipid phosphatase PTEN is also enriched in CCPs (Rosselli-Murai *et al*., 2018). Collectively, these studies raise the question of whether other components of the PI3K-Akt pathway may also localized with CCPs, and what additional proteins are required to support this signal recruitment and enrichment within plasma membrane clathrin structures.

The enrichment of PI3K has been reported in endosomes (Thapa *et al*., 2024), focal adhesions (Wang *et al*., 2024), and membrane rafts (Cizmecioglu *et al*., 2016). Notably, there is some overlap of CCPs and membrane raft components, such as following EGF stimulation (Puri *et al*., 2005). Further, the localization of PI3K may differ upon growth factor stimulation, as the enrichment of PI3K-Akt signals to focal adhesions was reported under basal conditions in the absence of growth factor stimulation (Wang *et al*., 2024). These studies collectively highlight the importance of nanoscale spatial organization of PI3K-Akt signaling and suggest that this spatial organization may be distinct in specific signaling contexts. The CCP enrichment of Gab1 (Garay *et al*., 2015; Lucarelli *et al*., 2017), the essential adaptor protein for PI3K recruitment and activation downstream of ligand binding by EGFR (Mattoon *et al*., 2004), suggests that PI3K itself could be enriched within CCPs, although this has not been examined.

Candidate proteins that may participate in the recruitment and regulation of PI3K-Akt signalling by EGFR within CCPs could be expected to associate with these signals. Nck1 and Nck2 (Non-catalytic region of tyrosine kinase) are adaptor proteins that each harbor three Src-homology 3 (SH3) domains and one Src-homology 2 (SH2) domain. Expressed across various tissues, Nck1/2 proteins boast over 200 binding partners and were previously considered functionally redundant (Brown *et al*., 2005). However, despite some overlapping roles, Nck1 and Nck2 exhibit non-redundancy in certain functions (Ngoenkam *et al*., 2014). The importance of Ncks results from interactions such as with EGFR, PAK1, SOS, and DOCK1 (Bywaters and Rivera, 2021). Ncks have been shown to increase migration and invasion of breast cancer cells (Morris *et al*., 2017) and play essential roles in mammary gland physiology (Golding *et al*., 2023).

Nck1/2 have been found to interact with CCP localized proteins dynamin, FCHSD2, intersectin, synaptojanin, and Ack1 (Wunderlich *et al*., 1999a; Jacquet *et al*., 2018). Moreover, Nck1/2 also associate with EGFR (Hake *et al*., 2008) and components of PI3K-Akt signaling including Gab1 (Abella *et al*., 2010; Leung *et al*., 2014). This raises the interesting possibility that Nck1/2 adaptors may serve as a link to coordinate clathrin-dependent PI3K-Akt signaling downstream of EGFR, which is the focus of this study. We first examine the localization of Nck1 and Nck2 to CCPs, finding that both Nck adaptors are enriched within CCPs. Nck2 participates in CCP initiation, assembly, and dynamics. Using siRNA gene silencing, we find that Nck1 and Nck2 are each required for phosphorylation of Akt upon EGF stimulation and may also negatively regulate EGF-stimulated EGFR phosphorylation. By tracking fluorescently-tagged PI3K expressed at controlled levels or endogenous PI3K, we find enrichment of PI3K within plasma-membrane clathrin structures upon EGF stimulation, which was dependent on Nck1/2 and Gab1. We also find essential roles for Nck1/2 in supporting cell viability and/or proliferation. Collectively, this work suggests partially non-redundant roles for Nck1 and Nck2 adaptor proteins in regulating CCP dynamics and EGFR signaling leading to PI3K-Akt activation, impacting cell physiology.

## Results

### Distinct localization of Nck1 and Nck2 in clathrin-coated pits

We first aimed to examine how Nck adaptor proteins may localize to CCPs and how they may regulate these structures. We are not aware of antibodies that allow specific and efficient immunofluorescence labelling of endogenous Nck1 or Nck2. To determine the localization of Nck adaptor proteins in ARPE-19 cells, we created stable cell lines that allow doxycycline-inducible expression of either Nck1-eGFP or Nck2-eGFP using the Sleeping Beauty transposon system (Kowarz *et al*., 2015; Zak and Antonescu, 2023). These cell lines allow titration of doxycycline for induction of eGFP-tagged Nck proteins at controlled levels of expression that approximate the expression of endogenous Ncks (Zak and Antonescu, 2023) (**Figure S1**).

Following EGF stimulation and fixation, the cells were stained with antibodies to label clathrin heavy chain and then subjected to TIRFM to image the cell surface. Both Nck1-eGFP and Nck2-eGFP cells exhibited Nck localization to a range of structures at the cell surface (**Figure 1A-B, S2A-B**), including punctate structures and structures that appear to be focal adhesions. While some Nck-eGFP structures did not overlap with clathrin structures at the plasma membrane, there are also many clathrin structures also appear to contain Nck1-eGFP or Nck2-eGFP. To determine if this represents enrichment of recruitment of Nck1-eGFP or Nck2-eGFP to clathrin structures relative to the rest of the cell surface, images were analyzed using automated detection and analysis of clathrin-labelled structures (CLS) performed using a Gaussian-based modeling approach (Aguet *et al*., 2013), as used previously to quantify the specific recruitment of proteins to CLSs (Delos Santos *et al*., 2017; Lucarelli *et al*., 2017; Cabral-Dias *et al*., 2022; Abousawan *et al*., 2023; Rahmani *et al*., 2023; Sugiyama *et al*., 2023; Orofiamma *et al*., 2024). The term CLS refers to clathrin structures detected in fixed samples, as distinguishing genuine clathrin-coated pits (CCPs) from short-lived subthreshold clathrin structures requires live cell analysis (Aguet *et al*., 2013; Kadlecova *et al*., 2017). To allow measurement of the selective recruitment of GFP-tagged Ncks to CLSs, we also performed this analysis in images in which one of the channels had been rotated 180 degrees, thus scrambling the position of CLSs objects relative to Nck (eGFP) channel, allowing measurement of non-selective overlap of Nck-eGFP within CLSs, as we have done previously to measure the recruitment of specific proteins to CLSs (Cabral-Dias *et al*., 2022; Sugiyama *et al*., 2023). We termed this condition “randomized” detection (rando).

**Figure 1.**
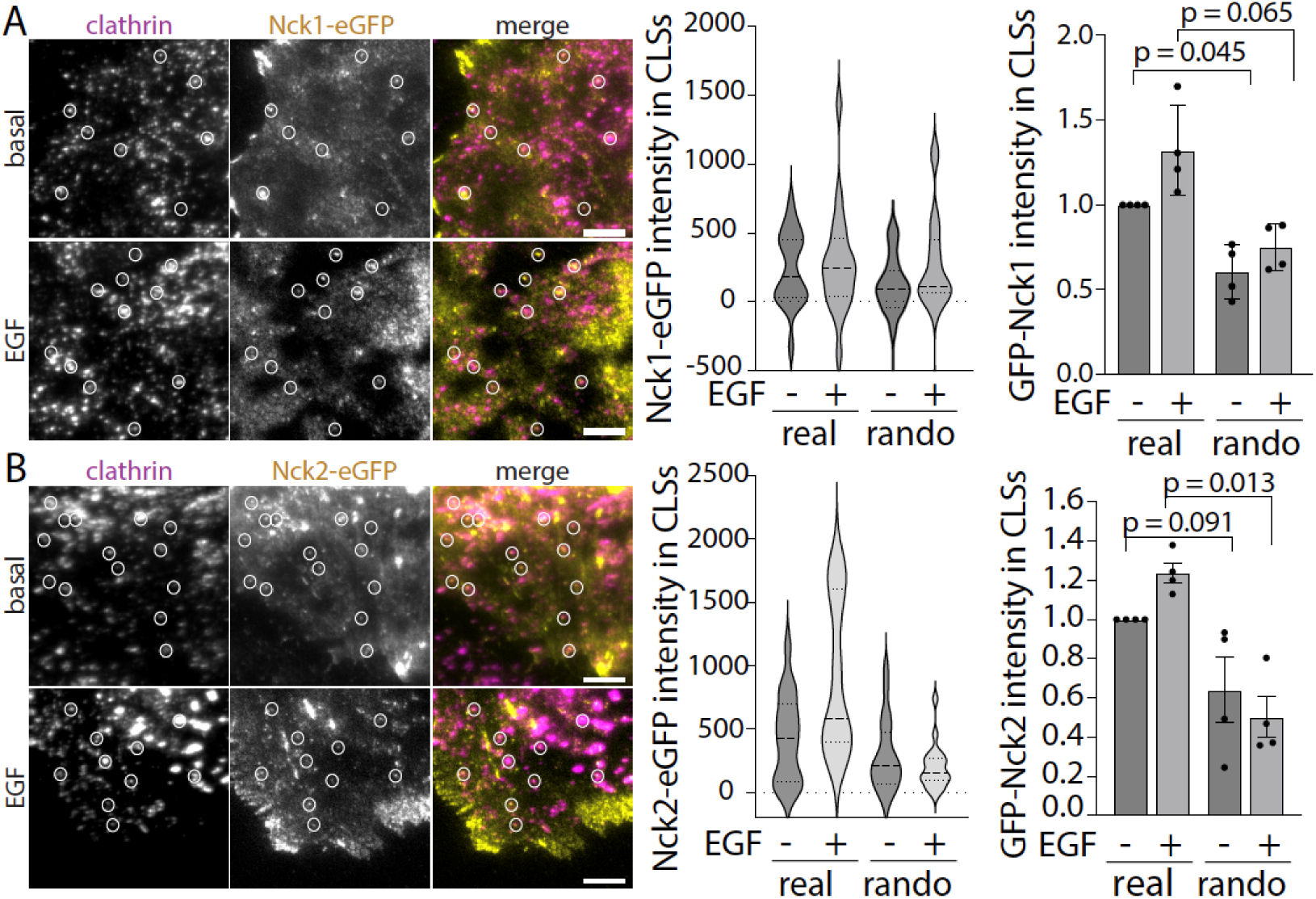
Nck1 and 2 are enriched in clathrin structures at the plasma membrane. ARPE-19 cells harboring stable transgenes for inducible expression of Nck1-eGFP or Nck2-eGFP were treated with 150ng/mL doxycycline for 24h. Then, on the day of the experiment, cells were stimulated with 20 ng/ml EGF for 5 min or left unstimulated (basal), then fixed and examined using TIRFM. Displayed (left) are representative images showcasing Nck1 or Nck2-associated clathrin structures. Scale = 5 µm. Full sized images shown in **Figure S1**. Images obtained by TIRFM underwent automated detection and analysis of CLSs, facilitating the quantification of Nck1-eGFP and clathrin in each identified structure. Displayed (middle) are the assessments of Nck1-eGFP or Nck2-eGFP fluorescence intensity within CLSs, illustrating the distribution of mean values from individual cells presented as a violin plot, with median (long dashed line) and 25th/75th percentiles (short dashed line) indicated. Also illustrated (right panels) are the fluorescence levels of Nck1-eGFP or Nck2-eGFP within CLSs from n = 4 independent experiments shown as mean ± SEM. Measurements are also included for image pairs where one channel was rotated 180° to randomize the positioning of Nck1 or Nck2 structures relative to clathrin structures (rando). The total number of CLSs quantified are as follows: Nck1-eGFP basal: 80, 4049; and EGF-stimulated: 80, 3338; Nck2-eGFP basal: 80, 4634, EGF-stimulated: 75, 5404. P-values shown determined by two-way ANOVA with a Tukey post-hoc test.

The presence of Nck1-eGFP or Nck2-eGFP within CLSs is shown for individual cells in a single experiment (**Figure 1 A-B**, middle panels), as well as the average results from multiple independent experiments (**Figure 1 A-B**, right panels). This analysis reveals that there is significant enrichment of Nck1-eGFP and Nck2-eGFP within CLSs compared to the randomized image pairs (**Figure 1A-B**). Upon stimulation with EGF at 20ng/mL for 5 minutes, the colocalization of Nck1-eGFP and Nck2-eGFP may increase within CLSs, although this was not statistically significant. That Nck1 and Nck2 each localize to CLS is consistent with their similar structure (Chen *et al*., 1998). This partial enrichment of Nck1/2 in plasma membrane clathrin structures, prompted us to examine their role in CCP initiation, assembly, and turnover.

### Nck2 selectively regulates clathrin-coated pit assembly and dynamics

To investigate the impact of Nck proteins on cellular dynamics, we used siRNA gene silencing to selectively suppress either Nck1 or Nck2 in ARPE-19 cells stably expressing eGFP fused to clathrin light chain (eGFP-CLC), followed by live-cell imaging using TIRFM. During imaging, cells were also stimulated with EGF. Interestingly, silencing Nck1 had no effect on the rate of nucleation of sub-threshold CLSs (sCLSs) (**Figure 2B**) or initiation of bona fide CCPs (**Figure 2C**). sCLSs are thought to represent transient stochastic assemblies of clathrin and other proteins at the cell surface, while CCP initiation represents the regulated start of CCP formation (Kadlecova *et al*., 2017). Moreover, silencing Nck1 had no discernable effect on CCP lifetimes, as observed by the mean CCP lifetimes (**Figure 2D**), by the proportion of short-lived CCPs that exhibit lifetimes <15s (**Figure 2E**), nor on or the proportion of persistent CCPs with lifetimes >300s (**Figure 2F**). This indicates that Nck1 contributes minimally to the regulation of CCP dynamics in ARPE-19 cells. Consistent with the lack of effect of Nck1 on CCP lifetimes and initiation rates, silencing Nck1 also did not impact the mean fluorescence intensity of eGFP-clathrin within CCPs (**Figure 2G**), indicating that silencing Nck1 does not impact the size of CCPs in ARPE-19 cells.

**Figure 2.**
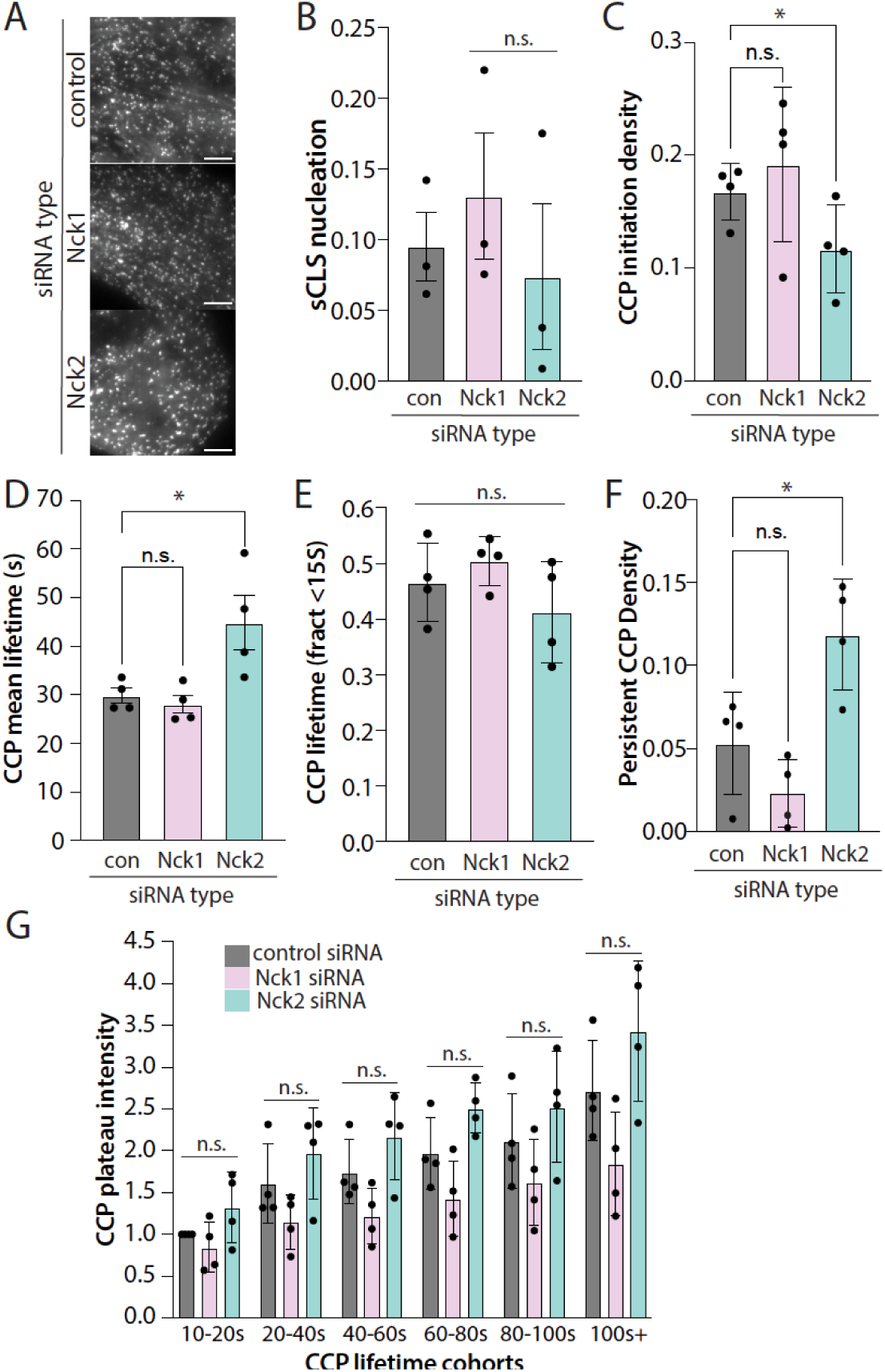
Nck2 silencing impacts CCP initiation and lifetimes. ARPE-19 cells stably expressing eGFP-clathrin light chain were treated with either Nck1 or Nck2 siRNA or with non-targeting siRNA (control). Following transfection, cells were subjected to imaging with near-simultaneous time-lapse TIRF microscopy while in being subject to stimulation with 5 ng/mL EGF. These time-lapse image series were subjected to automatic detection, tracking, analysis of clathrin structures as described in the *Methods* section. Shown are the nucleation rates of sCLSs **(B)** and initiation of CCPs **(C)**. Also shown are the intensities of eGFP-clathrin in CCPs based on mean lifetimes of CCPs **(D)** lifetime fractions of <15s **(E)**, persistent CCPs **(F)**, and intensity of eGFP-clathrin within clathrin structures shown as the ‘plateau intensity’ **(G)**. In each case the **(B-G)** the data is presented as the mean (bar) ± SEM; *n = 4* independent experiments, as well as individual values from each experiment (dots); ∗, *p* < 0.05 determined by one-way (B-F) or two-way (G) ANOVA with Holm-Šídák’s multiple comparisons test. The number of total CCP trajectories and cells (respectively) for each condition are: control 23662, 44; Nck1 siRNA 22268, 42; Nck2 siRNA 15810, 37 *, p < 0.05. abbreviations: CCPs, clathrin-coated pit; sCLS, subthreshold clathrin labeled structure; TIRF, total internal reflection fluorescence.

In contrast, while silencing Nck2 did not impact the rate of nucleation of sub-threshold CLSs (sCLSs) (**Figure 2B**), silencing Nck2 decreased the rate of initiation of bona fide CCPs (**Figure 2C**), consistent with a role for Nck2 in regulation of the early stages of protein assembly within CCPs. In addition, Nck2 silencing increased the mean lifetime of CCPs (**Figure 2D**). The latter could be the result of an increased proportion of early CCPs becoming stabilized into productive CCPs or a delay in the resolution of longer-lived CCPs, perhaps as a result of defects in scission from the plasma membrane. Interestingly, silencing Nck2 did not impact the proportion of short-lived CCPs with lifetimes <15s (**Figure 2E**) yet increased the proportion of long-lived persistent CCPs (**Figure 2F**). This lifetime analysis suggests that while Nck2 may be largely dispensable for stabilization of early CCPs, Nck2 may be required for efficient maturation of CCPs and/or for regulation of CCP scission from the plasma membrane, consistent with Nck2 interaction with the GTPase dynamin (Wunderlich *et al*., 1999b).

### Nck1 and Nck2 regulate PI3K-Akt signaling

Plasma membrane clathrin structures are important for the regulation of PI3K-Akt signaling by EGFR (Sigismund *et al*., 2008; Garay *et al*., 2015; Leyton-Puig *et al*., 2017; Rosselli-Murai *et al*., 2018; Pascolutti *et al*., 2019; Alfonzo-Méndez *et al*., 2022; Cabral-Dias *et al*., 2022). To determine the roles of Nck1 and Nck2 in EGFR signaling, we first investigated whether these adaptors contribute to the activation of the PI3K-Akt pathway. Specifically, we examined the effects of Nck1 or Nck2 silencing on Akt phosphorylation, using an antibody that recognizes the phosphorylated form of all Akt isoforms. Notably, EGF-stimulated Akt phosphorylation was markedly reduced in cells treated with siRNAs targeting either Nck1 or Nck2 (**Figure 3A**), suggesting non-redundant functions for Nck1 and Nck2 in EGF-stimulated Akt activation.

**Figure 3.**
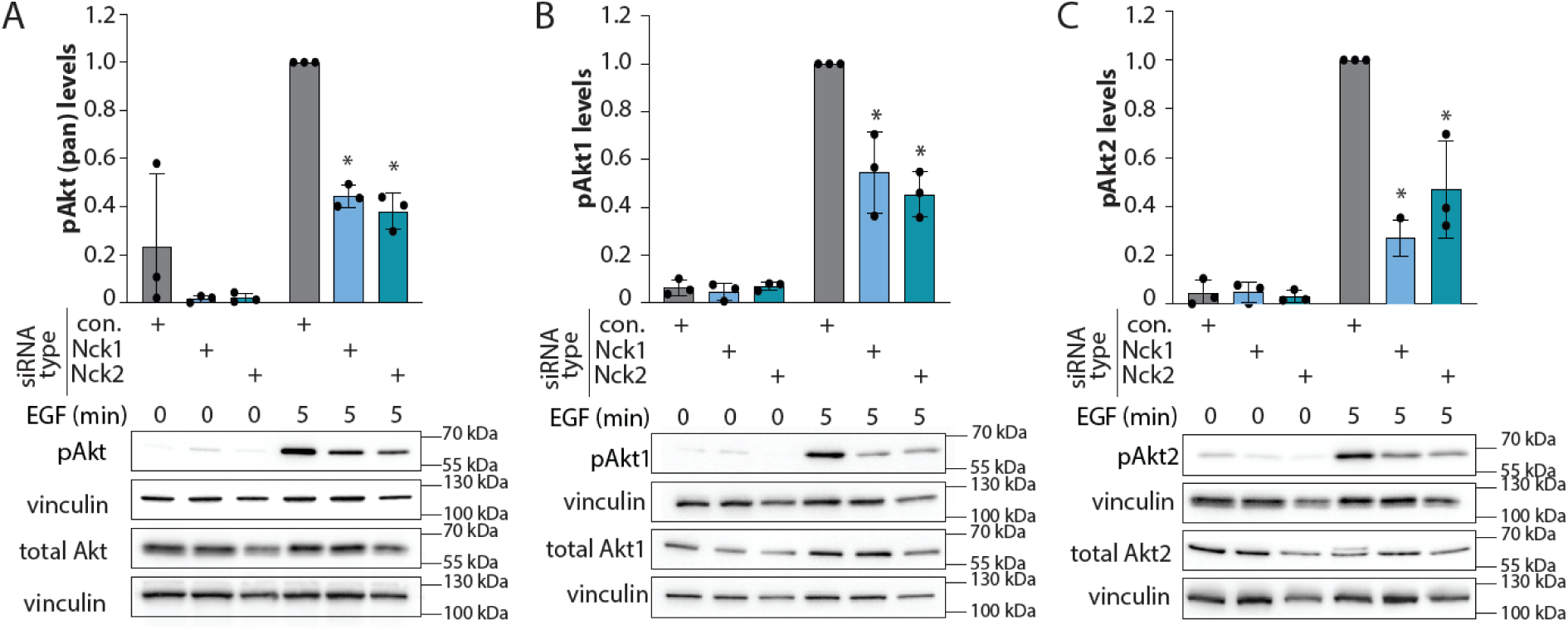
Nck1 and Nck2 each regulate EGF-stimulated Akt phosphorylation. ARPE-19 cells were transfected with siRNA targeting Nck1, Nck2, or nontargeting siRNA (control), followed by stimulation with 5ng/ml EGF for 5 min. **(A)** Western blotting of whole-cell lysates probed with anti-phospho-Akt (Ser473), anti-phospho-Akt1 or anti-phospho-Akt2. **(B)** Also shown are the mean ± SEM anti-phospho-Akt (Ser473), anti-phospho-Akt1 or anti-phospho-Akt2 with points representing individual experiment measurements; n = 3; *, P < 0.05 determined by two-way ANOVA with Tukey post-hoc test, relative to the control siRNA-treated EGF-stimulated condition.

We next examined the mechanisms by which Nck1 and Nck2 contribute to Akt phosphorylation by scrutinizing the effects of Nck1 and Nck2 silencing on phosphorylation of individual Akt isoforms. Akt1 is activated in response to membrane recruitment by binding to phosphatidylinositol-3,4,5-triphosphate (PIP_3_), while Akt2 is activated by binding to PI(3,4)P_2_, a lipid derived from PIP_3_ through the action of the lipid phosphatase SHIP2 (Goulden *et al*., 2018; Liu *et al*., 2018). Silencing Nck1 or Nck2 individually each led to a decrease in EGF-stimulated phosphorylation of both Akt1 (**Figure 3B**) and Akt2 (**Figure 3C**). Given that PIP_3_ production is crucial for both Akt isoforms, these findings suggest that Nck1 and Nck2 may regulate PI3K and PIP_3_ production upon EGF stimulation.

Notably, we also observed similar results with distinct siRNA sequences targeting Nck1 and Nck2 in ARPE-19 cells (**Figure S4**), illustrating that the effects of silencing each Nck adaptor protein is likely to be specific. Moreover, we also examined the impact of silencing Nck1 and Nck2 in another cell line, the breast cancer cell line MDA-MB-231, observing similar results showing that silencing of either Nck1 or Nck2 resulted in a loss of EGF-stimulated Akt phosphorylation without impairing other aspects of EGFR signaling (**Figure S5**). These results show that Nck1 and Nck2 specifically regulate EGFR signaling leading to Akt phosphorylation.

To determine whether the loss of Akt phosphorylation upon EGF stimulation was due to changes in EGFR activation or downstream signaling, we analyzed EGF-stimulated EGFR phosphorylation following Nck1 or Nck2 silencing. Silencing of Nck1 or Nck2 resulted in an increase in EGF-stimulated EGFR phosphorylation (**Figure 4A**). This indicates that the loss of EGF-stimulated Akt phosphorylation was not due to a reduction of cell surface EGFR or a defect in EGF-stimulated activation of EGFR activity. Consistent with Nck1 or Nck2 silencing supporting at least normal EGF-stimulated EGFR phosphorylation, there was no significant effect on EGF-stimulated Erk phosphorylation upon silencing of Nck1 or Nck2 (**Figure 4B**). Taken together, these results suggest that Nck1 and Nck2 each modulate EGFR signaling downstream EGFR binding and autophosphorylation, leading to selective regulation of activation of the PI3K-Akt signaling pathway.

**Figure 4.**
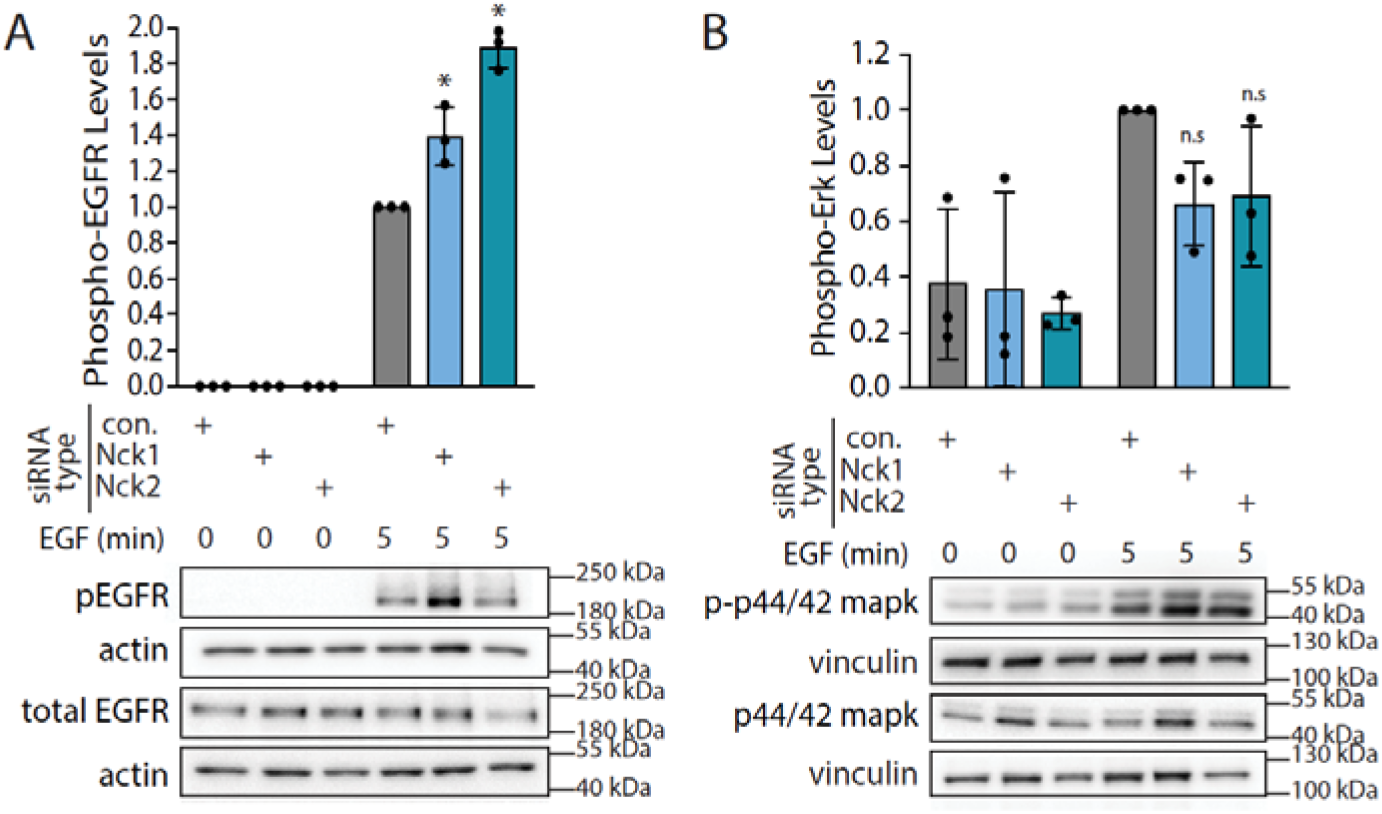
Nck1 or Nck2 silencing increases EGF-stimulated EGFR phosphorylation but does not affect the MAPK pathway. ARPE-19 cells were transfected with siRNA targeting Nck1, Nck2, or nontargeting siRNA (control), followed by stimulation with 5 ng/ml EGF for 5 min. **(A)** Western blotting of whole-cell lysates probed using anti-phospho-EGFR (pY1068); also shown are the mean ± SEM phospho-EGFR with points representing individual experiment measurements; *n = 3*; *, P < 0.05, relative to the control siRNA-treated EGF-stimulated condition, determined by two-way ANOVA with Tukey post-hoc test. **(B)** Western blotting of whole-cell lysates probed with anti-phospho-Erk; also shown are the mean ± SEM phosphor-Erk levels with points representing individual experiment measurements; *n = 3*.

### Nck1 and Nck2 regulate PI3K signaling complex recruitment to CCPs

Nck1 and Nck2 both support activation of PI3K-Akt signaling upon EGF stimulation (**Figure 3**), and plasma membrane clathrin structures regulate this signaling pathway (Sigismund *et al*., 2008; Garay *et al*., 2015; Leyton-Puig *et al*., 2017; Rosselli-Murai *et al*., 2018; Pascolutti *et al*., 2019; Alfonzo-Méndez *et al*., 2022; Cabral-Dias *et al*., 2022). Thus, we next sought to examine how Nck adaptors may regulate the localization of PI3K-Akt signals. To do so, we examined the localization of PI3K itself within the plasma membrane relative to plasma membrane clathrin structures, using antibodies to detect the endogenous p110β catalytic subunit (Thapa *et al*., 2020) and TIRFM in cells expressing eGFP-clathrin (**Figure 5A**). As for the analysis shown in Figure 1, we performed automated quantification of the enrichment of PI3K p110β in CLS in real image pairs, as well as in the same image datasets in which the PI3K p110β image had been rotated 180 degrees to randomize the spatial relationship between the two markers (randomized). This revealed that EGF stimulation elicits selective enrichment of PI3K p110β to CLS within these TIRF images (**Figure 5A**, right panels).

**Figure 5.**
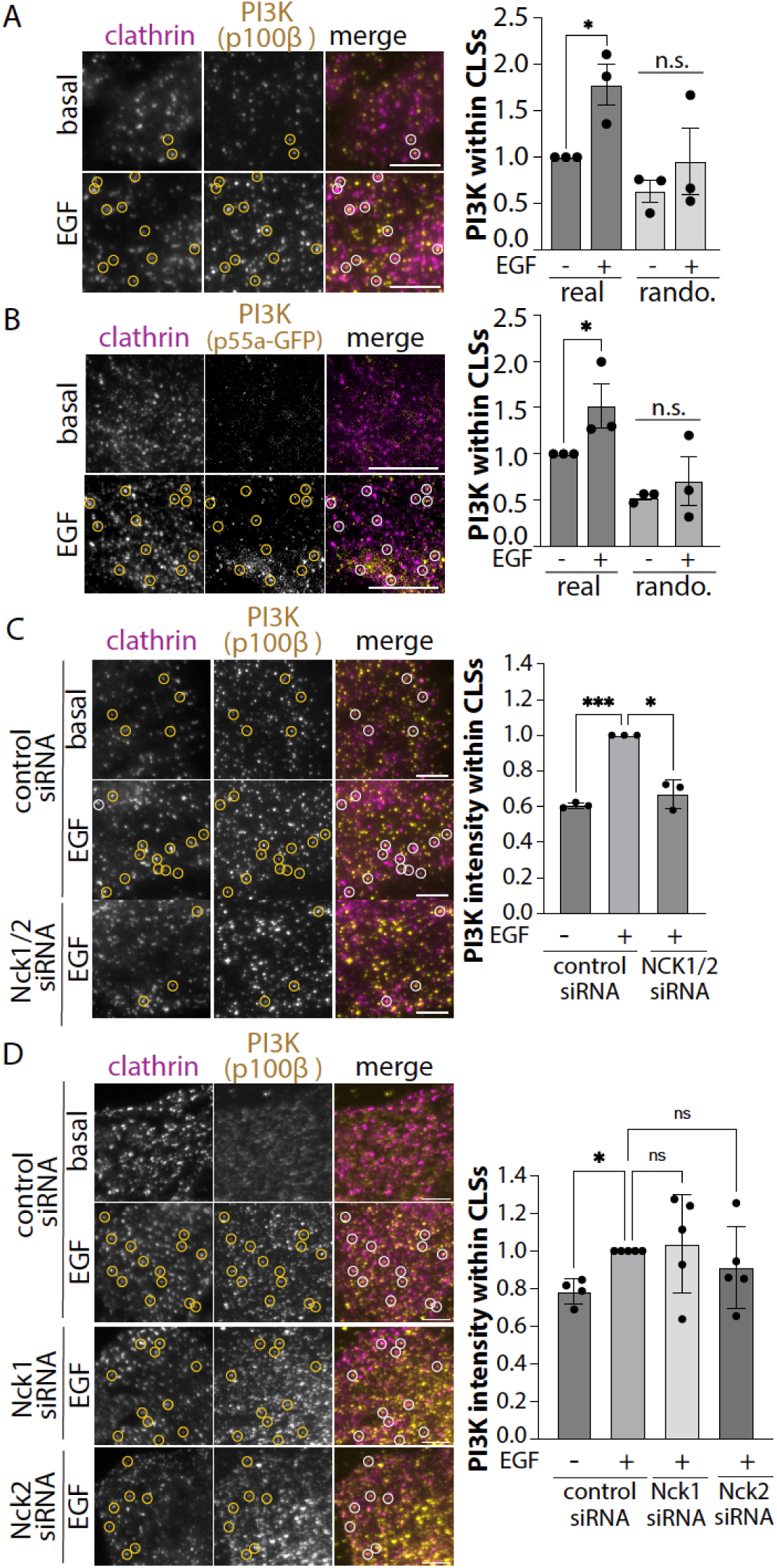
Nck1/2 regulate PI3K recruitment to plasma membrane clathrin structures. ARPE19 cells stably expressing eGFP-clathrin light chain (***A, C-D***) or harboring a transgene for inducible expression of eGFP-p55α (regulatory PI3K subunit) and subjected to 24h treatment with 150 nM doxycycline (***B***) were used in these experiments. In C-D, cells were also subject to siRNA silencing as indicated. Following stimulation with 20 ng/mL EGF as indicated, cells were fixed and stained with antibodies to detect clathrin heavy chain (A, C-D) or p110β (PI3K catalytic subunit); samples were then examined using TIRFM. Displayed (left) are representative images showing PI3K-associated clathrin structures. Scale = 5 µm. Images obtained by TIRFM underwent automated detection and analysis of CLSs, facilitating the quantification of PI3K subunit enrichment within each identified clathrin structure. Shown are the fluorescence levels of PI3K subunits within CLSs from n = 3 (A-C) or 5 (D) independent experiments shown as mean ± SEM. Measurements are also included for image pairs where one channel was rotated 180° to randomize the positioning of PI3K relative to clathrin structures (rando). The total number of CLSs and cells (respectively) quantified are as follows: (A) basal: 6228, 71, and EGF-stimulated: 5243, 80; (B) basal: 7852, 28, and EGF-stimulated: 10702, 37; (C) control siRNA basal: 15147, 75; control siRNA EGF-stimulated: 17264, 72; Nck1/2 siRNA EGF-stimulated: 17596, 68; (D) control siRNA basal: 30751, 100; control siRNA EGF-stimulated: 26344, 100; Nck1 siRNA EGF-stimulated: 25254, 100; Nck1 siRNA EGF-stimulated: 25384, 100; *, p < 0.05 determined by Two-way (A-B) or one-way (C-D) ANOVA with Tukey post-hoc test.

To support the detection of PI3K p110β in these experiments, we examined the effect of silencing of Gab1, an essential adaptor protein for plasma membrane recruitment of PI3K upon EGF stimulation (Mattoon *et al*., 2004) that we also previously showed to be enriched in plasma membrane clathrin structures (Garay *et al*., 2015; Lucarelli *et al*., 2016, 2017). Silencing Gab1 led to a loss of PI3K p110β detection within plasma membrane CLSs (**Figure S6A**), consistent with specific detection of PI3K p110β within these structures. To further complement these observations, we also generated a stable cell line that allows expression of an eGFP-tagged p55α regulatory subunit of PI3K (**Figure 5B**). Consistent with our observations with detection of endogenous PI3K p110β, EGF stimulation triggered a selective increase in the recruitment of eGFP-p55a to CLS at the cell surface (**Figure 5B**, right panels).

We next sought to resolve how Nck proteins may regulate PI3K recruitment to CCPs. Silencing of both Nck1 and Nck2 led to a reduction of PI3K p110β recruited to plasma membrane CLSs upon EGF stimulation (**Figure 5C**). Interestingly, silencing of each of Nck1 and Nck2 individually did not significantly impact the EGF-stimulated gain in enrichment of PI3K p110β within CLSs (**Figure 5D**), suggesting that Nck adaptors were at least partially redundant for recruitment of PI3K to plasma membrane clathrin structures.

To resolve how Ncks may control the recruitment of PI3K to plasma membrane clathrin structures, we examined how Ncks impact the recruitment of ligand-bound EGFR to these structures. We treated ARPE-19 cells expressing eGFP-clathrin within rhodamine-EGF for 5 min, followed by fixation, imaging by TIRFM, and detection of ligand-bound EGFRs within plasma membrane clathrin structures (**Figure 6**). These experiments revealed that silencing either Nck1 or Nck2 alone was sufficient to significantly reduce the recruitment of EGFR to plasma membrane clathrin structures. Moreover, silencing Gab1 also reduced the recruitment of ligand-bound EGFR to CLSs (**Figure S6B**). Taken together with our observations that silencing Nck1 or Nck2 did not suppress EGFR phosphorylation and instead appears to even increase EGF-stimulated EGFR phosphorylation (**Figure 4A**), this suggests that Nck1 and Nck2 are each required to support the recruitment of ligand-bound, active EGFR to plasma membrane clathrin structures, along with recruitment of signaling complexes that drive activation of the PI3K-Akt pathway therein.

**Figure 6.**
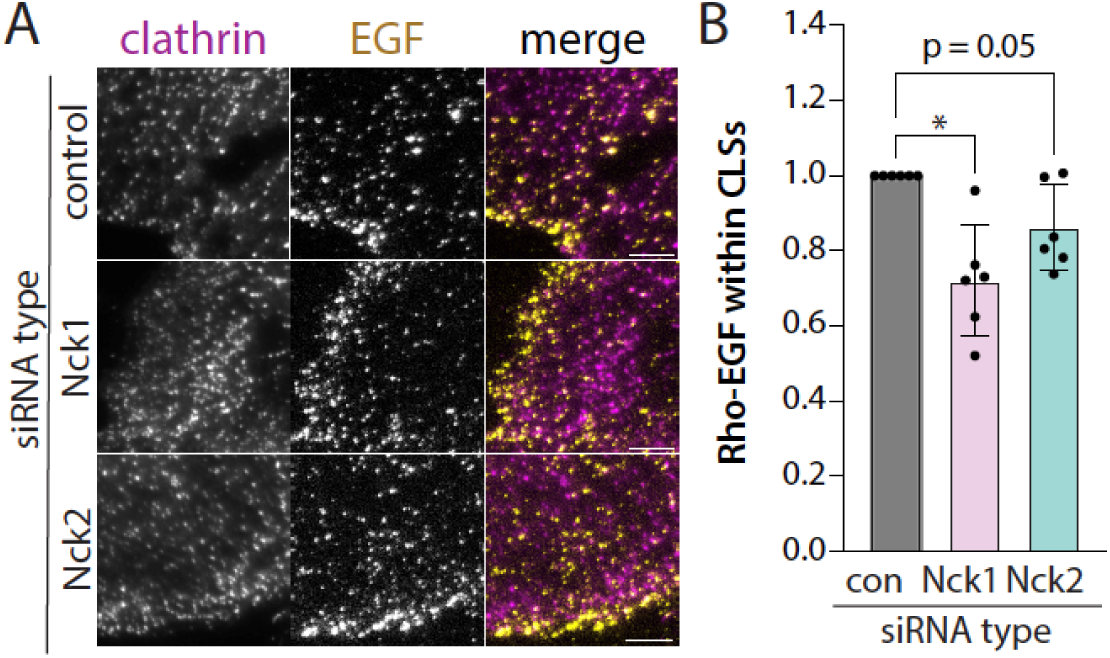
Nck1 and 2 each regulate EGFR recruitment to plasma membrane clathrin structures. ARPE19 cells were subject to siRNA silencing as indicated. Cells were stimulated with 20 ng/mL rhodamine-EGF as indicated, after which cells were fixed and then examined using TIRFM. Displayed (left) are representative images showing rhodamine-EGF-associated clathrin structures. Scale = 5 µm. Images obtained by TIRFM underwent automated detection and analysis of CLSs, facilitating the quantification of rhodamine-EGF enrichment within each identified structure. Shown are the fluorescence levels of rhodamine-EGF within CLSs from n = 3 independent experiments shown as mean ± SEM. The total number of CLSs and cells quantified (respectively) are as follows: control siRNA: 34598, 120, Nck1 siRNA: 30195, 120; Nck2 siRNA: 29450, 120, *, p < 0.05 determined by one-way ANOVA with Tukey post-hoc test.

### Nck1 and Nck2 regulate cell survival and/or proliferation

The PI3K-Akt pathway leads to the phosphorylation of several targets including some that regulate the mammalian target of rapamycin complex 1 (mTORC1), a key regulator of cellular metabolism, immune responses, proliferation, autophagy, and migration. Collectively, the targets of the PI3K-Akt pathway lead to enhanced cell proliferation and increased cell viability (Manning and Toker, 2017; Sugiyama *et al*., 2019; He *et al*., 2021). To investigate the role of Nck1 and Nck2 in modulating PI3K-Akt signaling and their effects on cellular proliferation and viability, we performed siRNA gene silencing of Nck1 and Nck2 in ARPE-19 cells and monitored cell proliferation using an automated imaging system to quantify cellular confluence over time.

Silencing each of Nck1 and Nck2 in ARPE-19 cells resulted in a time-dependent reduction in cell confluence compared cells treated with control siRNA (**Figure 7A-B**), with Nck2 silencing appearing to have a more pronounced effect. Treatment of these samples in parallel with CellTox-green allowed identification of non-viable cells during this experiment. Silencing of Nck1 or Nck2 increased the detection of non-viable cells (**Figure 7A, C**), with Nck2 silencing again having a more pronounced effect. These findings suggest that while perturbation of each of Nck1 or Nck2 reduces proliferation and/or survival rates, Nck2 may play a more critical role in supporting cell proliferation and/or survival in ARPE-19 cells. The significant impact of Nck2 silencing on proliferation and/or survival is consistent with the requirement for Nck2 in regulation of clathrin-coated pit dynamics in addition to the requirement of both Nck isoforms regulating the full activation of Akt and PI3K signaling upon EGF stimulation.

**Figure 7.**
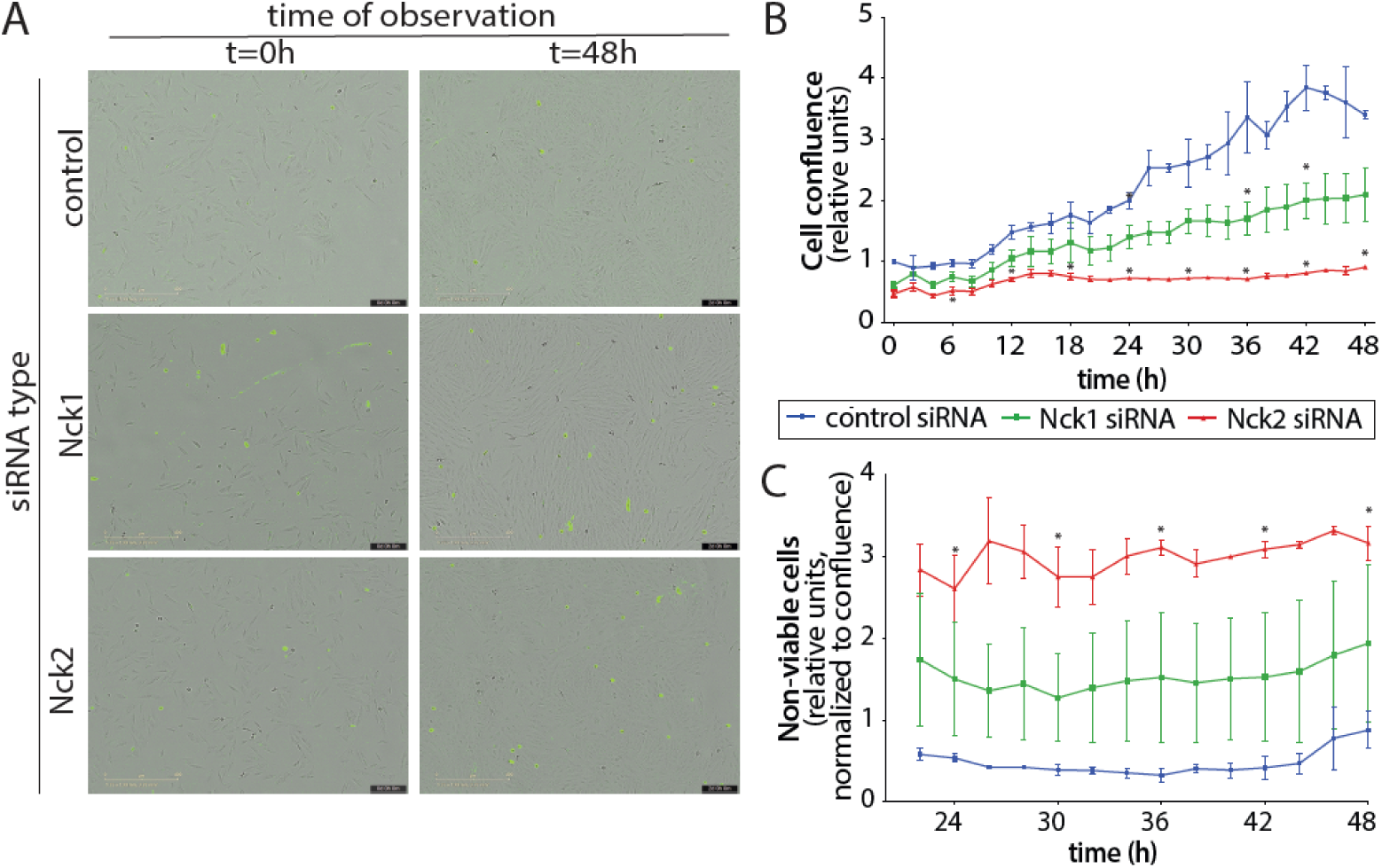
Nck1 and Nck2 silencing regulates cell proliferation and/or viability. ARPE-19 cells were subjected to Nck1 or Nck2 silencing, or treatment with non-targeting (control) siRNA. Following transfection, cells were subjected to time-lapse imaging using an Incucyte SX5 environmentally-controlled microscope for 48 h to measure cellular growth and viability. **(A)** Phase contrast images of ARPE-19 cells overlaid with CellTox green fluorescence signal at the indicated time in points (hours). Scale bars: 400µm. **(B)** Quantification of cellular confluence normalized to t = 0h in the control siRNA condition. **(C)** Quantification of cellular death labelled with CellTox at t = 12h to t = 48h. Data is presented as the mean ± SEM of *n*=3 independent experiments in **(C)** and **(D)**.

## Discussion

We examined the plasma membrane localization and contribution of Nck adaptor proteins to EGFR action leading to PI3K-Akt signaling and to clathrin-coated pit dynamics. Our results suggest that Ncks play a regulatory role in the formation of clathrin coated pits and in EGFR signaling, phenomena which are increasingly appreciated as intertwined.

### Mechanism of regulation of CCP dynamics by Ncks

Silencing Nck2 but not Nck1 impacted CCP dynamics in EGF-stimulated ARPE-19 cells (**Figure 2**). Both Nck1 and Nck2 were detected within plasma membrane clathrin structures over background (or the signal expected due to random overlap of Ncks and clathrin) (**Figure 1**). This result is consistent with Nck adaptor proteins being able to interact with clathrin structure localized proteins (Jacquet *et al*., 2018). Interestingly, in cells with arrested CCPs that result from knockout of all three dynamin isoforms, Nck1 was detected as one of the proteins with increased levels of phosphorylation (Shen *et al*., 2011), consistent with localization of Nck1 to plasma membrane clathrin structures.

We are not aware of any previous studies that have examined the role of Nck adaptor proteins in CCP dynamics. Silencing of Nck2 resulted in a reduction of CCP initiation, indicating that Nck2 contributes to early stages of CCP assembly that allow the few molecules of AP2 and clathrin to progress from largely stochastic nucleation of transient structures to a bona fide CCP (Kadlecova *et al*., 2017; Bhave *et al*., 2020). For CCPs that initiate, silencing Nck2 silencing did not significantly impact CCP size but instead increased mean CCP lifetimes (**Figure 2**). Taken together with the finding that Nck2 silencing did not impact the fraction of CCPs with lifetimes <15s (**Figure 2E**) and significantly increased the incidence of persistent CCPs (**Figure 2F**), this suggests that in addition to a role in early CCP assembly, Nck2 also has a role in late stages of CCPs, perhaps in promoting CCP scission.

Interestingly, work by Schmid & col. used a systematic approach to examine the functional contribution of 67 endocytic accessory proteins to CCP dynamics (Bhave *et al*., 2020). These proteins could be clustered based on the phenotype resulting from their loss-of-function. Several clusters recapitulated some of the phenotypes of Nck2 silencing, such as minimal effect on sCLS nucleation, a decrease in initiation of bona fide CCPs and an increase in CCP lifetimes. However, none of the clusters appeared to entirely recapitulate the effect of Nck2 silencing on increasing the fraction of persistent CCPs. Further, the ability of Nck proteins to interact with many endocytic accessory proteins (Jacquet *et al*., 2018) makes the model by which Nck proteins contribute to both early and late stages of CCP dynamics somewhat complex. Nonetheless, Nck1/2 also interact with the juxtamembrane domain of EGFR (Hake *et al*., 2008), and we observed that silencing of each of Nck1 or Nck2 suppressed the recruitment of EGFR to plasma membrane clathrin structures (**Figure 6**). Nck adaptors also bind to many other signaling receptors, including Nephrin (Martin *et al*., 2020), T-cell Receptor (Paensuwan *et al*., 2016), and B-cell Receptor (Castello *et al*., 2013), and to phospho-tyrosine containing motifs within the PDGF, VEGF, and Eph receptor tyrosine kinases (Alfaidi *et al*., 2021). A possible model by which Nck adaptors regulate the early stages of CCP formation including the initiation of bona fide CCPs is by regulating the recruitment of many receptors to CCPs, as receptor recruitment or clustering has been shown to promote CCP initiation (Liu *et al*., 2010). Indeed, such a role has been proposed for Nck adaptors in clustering Nephrin receptors that promotes Nephrin endocytosis (Martin *et al*., 2020).

The additional contribution of Nck2 to the late stages of CCP formation could be due to its role in remodeling the actin cytoskeleton. In mammalian cells, the actin cytoskeleton can contribute to regulation of CCP assembly, formation, and scission at multiple stages (Merrifield *et al*., 2005; Taylor *et al*., 2012; Grassart *et al*., 2014; Almeida-Souza *et al*., 2018; Yang *et al*., 2022), and the impact of the actin cytoskeleton on CCP may be context-dependent (Miya Fujimoto *et al*., 2000; Akamatsu *et al*., 2020). The domain architecture of Nck adaptors bears some similarity to other clathrin-localized adaptor proteins such as intersectin and CIN85, which have five and three tandem SH3 domains, respectively, and are CCP-localized proteins (Yamabhai *et al*., 1998; Kowanetz *et al*., 2003; Nikolaienko *et al*., 2009; Jin *et al*., 2024). From this emerges a possible model for Nck2 to regulate late stages of CCP formation by modulating actin remodeling related to vesicle scission. Ncks also interact with late-stage CCP proteins such as dynamin (Wunderlich *et al*., 1999a; Martin *et al*., 2020), suggesting another possible mechanism of regulation of CCP dynamics by Ncks in mediating CCP scission. However, while it is tempting to conclude that the regulation of CCP assembly by Nck2 results from localization of Nck2 to CCPs, it is also possible that this regulation is indirect, with Nck2 regulating CCP dynamics by control of cellular processes that subsequently impact CCP dynamics.

### Regulation of PI3K-Akt signaling by Ncks

The PI3K-Akt signaling pathway heavily relies on protein interactions mediated by SH2 and SH3 domains present in adaptor proteins. While SH2 interactions exhibit a relatively high affinity for pY motifs (e.g., Kd ∼30nM for Grb2’s SH2 domain binding to the motif surrounding pY1068 on EGFR (Batzer *et al*., 1994)), SH3 domain interactions with proline-rich domains (PRDs) are comparatively weaker, typically ranging from ∼10-100 µM Kd, and with a 30 µM Kd specifically between the SH3 of Grb2 and the proline-rich domain of Gab1 (Skolnik et al., 1993a; Batzer et al., 1994; Lewitzky et al., 2001; Harkiolaki et al., 2009). Moreover, Grb2’s SH3 domains also interact with other PRD-domain proteins like SOS and dynamin (Lewitzky *et al*., 2001). This relatively weak interaction of Grb2 with Gab1 suggests that signaling outcome and specificity isn’t solely governed by SH3 domains, indicating the involvement of additional regions within interacting proteins as has been proposed for SH3-dependent interactions (Kazlauskas *et al*., 2016; Dionne *et al*., 2021).

Within this context, Nck adaptors may link EGFR to a network of additional interactions, such as within plasma membrane clathrin structures, that support PI3K activation, leading to PIP_3_ production and Akt activation. In support of this model, we saw that silencing Ncks resulted in loss of recruitment of EGF-bound EGFR and PI3K to CCPs upon EGF stimulation, as well as a reduction in EGF-stimulated Akt phosphorylation. Since the loss of Ncks resulted in impaired phosphorylation of both Akt1 and Akt2 and given that PIP3 production is common to activation of both Akt isoforms (Liu *et al*., 2018), these findings suggest that Ncks act at or upstream of PIP_3_ production within EGFR signaling. Since there was no reduction of EGF-stimulated EGFR phosphorylation upon Nck silencing, this suggests that Ncks are dispensable for EGFR ligand binding, dimerization, and activation of the kinase domain.

By contributing additional, low affinity interactions, Ncks may enhance the interaction of signaling intermediates of the PI3K-Akt pathway associated with EGFR. At least some of these low affinity interactions may arise from an interaction network within CCPs. Consistent with this, we observed that silencing Nck1 or Nck2 resulted in a reduction of EGFR recruitment to CCPs (**Figure 6**) and that combined silencing of Nck1 and Nck2 resulted in a loss of EGF-stimulated enrichment of PI3K within CCPs. Interestingly, the SH2 domains of Nck1 and Nck2 each interact with Gab1 via motifs harboring a phosphorylated tyrosine residue(Leung *et al*., 2014), including pY407 on Gab1 (Abella *et al*., 2010). In addition to proline-rich domains, there are also other motifs that can bind SH3 domains (Teyra *et al*., 2017), suggesting that additional interactions of such motifs on Gab1 and SH3 domains on Ncks further strengthen signaling outcomes. In this context, Nck adaptors may function alongside Gab1 to strengthen a complex between EGFR, Gab1, and PI3K to promote activation of Akt, similarly to a role for Ncks in promoting the recruitment of the BCAP adaptor for PI3K activation in B-cell receptor signaling (Castello *et al*., 2013).

Nck proteins may have unique and non-redundant roles in regulating EGFR signaling, as silencing either of Nck1 and Nck2 led to defects in EGF-stimulated Akt phosphorylation (**Figure 3**), EGFR recruitment to CCPs (**Figure 6**), and reduced cell proliferation and viability in ARPE-19 cells (**Figure 7**). This is consistent with distinct, non-redundant roles in mammary gland development reported for Nck isoforms (Golding *et al*., 2023). However, Nck adaptors may have multiple functions in regulating PI3K-Akt signaling, in some of which Ncks may be functionally redundant, as silencing both Nck1 and Nck2 was required to elicit a significant loss in PI3K enrichment within CCPs in ARPE-19 cells (**Figure 5**). Nck1 and Nck2 differ most in sequence similarity in the linker region between the SH3 domains (Teyssier *et al*., 2024), suggesting that isoform-specific roles may be derived from the impact of this region. Collectively, these results suggest that Nck adaptors have several functions impacting activation of PI3K-Akt signaling.

### Clathrin Coated pits, Ncks, and signaling

The localization of Nck2 to clathrin structures and its role in regulating CCP dynamics, together with roles for Nck1 and Nck2 in modulating signaling intermediate recruitment to CCPs and Akt signaling suggest a link between plasma membrane clathrin structures and PI3K-Akt signaling. We have previously shown that clathrin is essential for EGF-stimulated activation of Akt (Garay *et al*., 2015). Since inhibition of vesicle formation by targeting dynamin did not impact EGF-stimulated Akt activation, we proposed that plasma membrane clathrin structures, rather than vesicle formation, are required to scaffold PI3K-Akt signaling at the plasma membrane. Consistent with this, we and others have identified Gab1 (Garay *et al*., 2015; Lucarelli *et al*., 2016, 2017) as well as signaling terminators PTEN (Rosselli-Murai *et al*., 2018) within CCPs, leading to regulation of PI3K-Akt signaling. We also recently reported that CCP-localized recruitment of the Src-family kinase Fyn by the clathrin-binding adaptor TOM1L1 regulates the clathrin localization of the lipid phosphatase SHIP2, selectively required for the EGF-stimulated phosphorylation of Akt2 (Cabral-Dias *et al*., 2022).

This work builds on the model of enrichment of PI3K-Akt intermediates within CCPs by showing that PI3K is also enriched within CCPs following EGF stimulation. EGF stimulation elicited an increase in PI3K in plasma membrane clathrin structures in paired images, but not in randomized images. Since the PI3K detection in objects within randomized image pairs represents “random” sampling of the plasma membrane, the lack of significant PI3K enrichment in these randomized pairs suggest that clathrin structures are a major site of PI3K recruitment following EGF stimulation. Given that CCPs are at the interface of the plasma membrane and intracellular vesicles, these findings complement reports of PI3K localization to intracellular compartments (Thapa *et al*., 2020, 2024). Our observation of the localization of PI3K to CCPs at the plasma membrane to elicit PI3K-Akt signaling is consistent with observations of normal EGF-stimulated PI3K-Akt signaling in cells with knockout of all three dynamin isoforms (Sousa *et al*., 2012) or with dynamin perturbation by siRNA silencing or drug treatment (Garay *et al*., 2015). While beyond the scope of this study, future work aimed at dissecting distinct aspects of Nck function that may separately drive Nck recruitment to CCPs and interaction with Gab1 or other components of PI3K signaling will be very useful in defining how clathrin enrichment of signals regulates EGF-stimulated PI3K activation and PIP3 production.

In summary, this study reveals novel, partially non-redundant roles for Nck adaptor proteins in regulating EGFR signaling leading to activation of the PI3K-Akt pathway, and a novel role for Nck adaptors in regulating clathrin-coated pit formation. These findings contribute to a growing understanding of the interrelatedness and interdependence of signaling and endocytosis molecular machineries, particularly those involving plasma membrane clathrin structures.

## Acknowledgements

This work was supported by a Project Grant from the Canadian Institutes of Health Research to C.N.A. (186250).

## Materials and Methods

### Materials

Antibodies recognizing specific proteins were as follows (with species and catalog numbers indicated for each): Nck1 (15B9, rabbit monoclonal, 2319), phospho-EGFR (pY1068, rabbit monoclonal, 3777), phospho-Gab1 (pY627, rabbit monoclonal, 3233), pAkt1 (S473, rabbit monoclonal, 9018), pAkt2 (S474, rabbit monoclonal, 8599), total Akt (pan isoform, mouse, 2920), actin (D18C11, rabbit monoclonal, 8456T), GFP (D5.1 XP, rabbit monoclonal, 2956S) and EGF Receptor (D38B1 XP, rabbit monoclonal, 4267S) obtained from Cell Signaling Technology; phospho-Akt (pS473, pan isoform for Western blotting, 44-621G) obtained from Life Technologies; Nck2 (EPR3333, rabbit monoclonal, ab109239) obtained from Abcam; and anti-clathrin heavy chain (TD.1) used for immunoblotting obtained from Santa Cruz Biotechnology.

Fluorophore-conjugated or HRP secondary antibodies were from Jackson ImmunoResearch. Rhodamine-EGF was generated in-house as previously described (Lucarelli *et al*., 2017).

Wild-type (WT) human retinal pigment epithelial cells (ARPE-19; RPE cells herein) and ARPE-19 cells stably expressing CLC fused to enhanced GFP (RPE-eGFP-CLC) were previously described (Bone *et al*., 2017; Cabral-Dias *et al*., 2022; Abousawan *et al*., 2023; Rahmani *et al*., 2023). Cells were cultured in DMEM/F12 (Life Technologies) supplemented with 10% fetal bovine serum (Life Technologies), 100 U/ml penicillin, and 100 μg/ml streptomycin (Life Technologies) at 37°C and 5% CO_2_. MDA-MB-231 cell lines were maintained using RPMI (Sigma Aldrich) containing 10% fetal bovine serum (Life Technologies), 100 μg/ml streptomycin (Life Technologies) and 100 U/ml penicillin (Life Technologies) 37°C and 5% CO2.

### Generation of ARPE-19-NCK1 and ARPE-19-NCK2 Stable Cell Lines

pSBtet-BP was a gift from Eric Kowarz (Goethe-University of Frankfurt, Frankfurt, Germany, plasmid 60496; Addgene; http://n2t.net/addgene:60496;RRID:Addgene_60496; (Kowarz *et al*., 2015)). pCMV(CAT)T7-SB100 was a gift from Zsuzsanna Izsvak (MaxDelbrückCenter for Molecular Medicine, Berlin-Buch, Germany, plasmid 34879; http://n2t.net/addgene:34879; Addgene; RRID:Addgene_34879; (Mátés *et al*., 2009)).

For the generation of pSBtet-BP-NCK1-eGFP expression plasmid, an oligonucleotide encoding Nck1 fused to eGFP fused was generated by BioBasic, using the ORF sequence of Nck1 (NM_006153.6), followed by the sequence encoding a spacer peptide (GGG GGG TCT GGT GGC AGT GGA GGG GGA TCC), followed by the ORF for eGFP. This Nck1-spacer-eGFP synthesized oligonucleotide was cloned into the ORF of pSBtet-BP plasmid. A similar strategy was used to generate a plasmid encoding NCK2-eGFP, using the ORF from Nck2 (NM_001004720.3) instead. The pSBtet-BP plasmid encoding the p55α regulatory subunit of PI3K was generated using a sequence encoding eGFP, followed by a linker sequence (GGG GGG TCT GGT GGC AGT GGA GGG GGA TCC), followed by the ORF sequence of NM_181504.2.

Complete protocol and transfection method was used as previously described (Zak and Antonescu, 2023). Briefly, pSBtet-BP-NCK1eGFP, pSBtet-BP-NCK2-eGFP, or pSBtet-BP-eGFP-p55α were co-transfected along with pCMV(CAT)T7-SB100 using FuGene transfection reagent (Promega) into wild-type ARPE-19 cells. Starting 48h after transfection, cells were treated in 1 µg/mL puromycin for at least 2-3 weeks to select for stably transfected cells.

### siRNA transfections

siRNA transfections were performed using custom-synthesized siRNAs (Horizon Discovery) using RNAiMAX transfection reagent (Thermo Fisher Scientific) as per manufacturer’s instructions, as we have done previously (Cabral-Dias *et al*., 2022; Abousawan *et al*., 2023; Rahmani *et al*., 2023; Sugiyama *et al*., 2023). Cells were seeded on a 6-well plate 24 h before transfection. Briefly, cells were initially washed with phosphate-buffered saline (PBS) to remove FBS-containing growth media, followed by addition of OptiMEM media (Thermo Fisher Scientific). Cells were incubated with customized siRNA sequences at a cellular medium concentration of 220 pmol/L in OptiMEM media and incubated for 4 h, after which cells were washed and media replaced with regular growth medium. siRNA transfections were performed twice, 72 h and 48 h before each experiment. Sequences used were as follows: control (nontargeting, sense), CGU ACU GCU UGC GAU ACG GUU, (antisense) CCG UAU CGC AAG CAG UAC GUU; Nck1-1 (sense) UGA GAG AGA GGA UGA AUU AUU, (antisense) UAA UUC AUC CUC UCU CUC AUU; Nck2-1 (sense) GGA AGA ACA GCC UGA AGA AUU, (antisense) UUC UUC AGG CUG UUC UUC CUU, Nck1-2 (sense) GGA CAA AGG UGA UCG UCA UUU, (anti-sense) AUG ACG AUC ACC UUU GUC CUU; Nck2-2 (sense) CAG AAG AAG UUA UUG UGA UUU, (antisense) AUC ACA AUA ACU UCU UCU GUU. Unless otherwise specified, Nck1-1 and Nck2-1 sequences were used when referring simply to Nck1 and Nck2 siRNA.

### Whole-cell lysates and immunoblotting

This was performed as previously described (Delos Santos *et al*., 2017; Lucarelli *et al*., 2017; Abousawan *et al*., 2023). After transfection, and/or stimulation with EGF, whole-cell lysates were prepared in Laemmli sample buffer (0.5 M Tris, pH 6.8, glycerol, 10% SDS, 10% β-mercaptoethanol, and 5% bromophenol blue; all from Bio-Shop) supplemented with a protease and phosphatase inhibitor cocktail (1 mM sodium orthovanadate, 10 nM okadaic acid, and 20 nM protease inhibitor cocktail, each obtained from BioShop). Lysates were then heated at 65°C for 15 min and passed through a 27.5-gauge syringe. Proteins were resolved by glycine-Tris SDS-PAGE followed by transfer onto a polyvinylidene fluoride membrane; they were washed, blocked, and incubated with antibodies as previously described (Antonescu et al., 2011). Molecular weight markers used were Novex Sharp Pre-Stained Protein Standard (LC5800) or PageRuler Prestained Protein Ladder (26617; Thermo Fisher Scientific).

Images were obtained using a Bio-Rad ChemiDoc Touch Imaging System upon soaking membranes in Luminata Crescendo HRP substrate (Millipore Sigma). Typical exposure times varied between 1 and 60 s and were selected to ensure that signal was not saturated at any pixel. Images were quantified using ImageJ software (National Institutes of Health; Schneider et al.,2012) by signal integration in an area corresponding to the appropriate lane and band for each condition. This measurement was then normalized to the loading control (e.g., vinculin) signal and then normalized to the total protein signal, in some cases obtained after reblotting. In each experiment, the resulting normalized phospho-protein/total-protein signal in each condition was expressed normalized to the control siRNA condition stimulated with EGF for 5 min. Statistical analysis was performed with ANOVA followed by Tukey post-test, with P < 0.05 used as a threshold for establishing differences between experimental conditions.

### EGF Stimulation

All cells were serum deprived for 1 h before experimental assays unless otherwise stated, cells were stimulated with 5ng/mL of EGF for 5 min. Experiments involving rhodamine EGF were subjected to 1 h serum starvation, stimulated with 20ng/mL rhodamine-EGF for 5 min prior to being fixed.

### Fluorescence labeling and microscopy

For detection of cellular proteins and fluorescent EGF, indicated cells were fixed with 4% PFA for 30 min, following by quenching of fixative in 100mM glycine, cell permeabilization in 0.1% Trition X-100 (all solutions made in PBS), and blocking in Superblock Blocking Buffer (Thermo Fisher Scientific). Cells were then stained with primary for 1 h at room temperature and followed by incubation with appropriate corresponding secondary antibodies. Cells were maintained in PBS at 4°C prior to TIRF-M imaging.

Cell samples were imaged using a Quorum (Guelph, ON, Canada) Diskovery instrument, comprised Leica DMi8 microscope operating in TIRF mode, equipped with a 63×/1.49 NA TIRF objective with a 1.8× camera relay (total magnification 108×). Images from samples were acquired using a Zyla 4.2 megapixel sCMOS camera with using 488 nm, 561, or 627 nm laser illumination and 525/50, 620/60, or 700/75 emission filters. Fixed-cell TIRFM imaging was done at room temperature with samples mounted in PBS. For live-cell imaging experiments, cells were maintained at constant 37°C during imaging, in phenol-free DMEM/F12 media (Gibco) supplemented with 20 mM HEPES and 20 ng/mL EGF, with imaging at rate of 1 frame per 15 s for 5 min.

### TIRF microscopy image analysis of Clathrin, Nck1, Nck2 and EGFR structures

Detection and analysis of Clathrin, Nck1, Nck2 or rhodamine-EGF structures at the diffraction limit were conducted using methodologies previously established (Delos Santos *et al*., 2017; Lucarelli *et al*., 2017; Cabral-Dias *et al*., 2022; Sugiyama *et al*., 2023). Custom software developed in Matlab (Mathworks Corporation, Natick, MA) facilitated this analysis, following the protocols outlined in (Aguet *et al*., 2013). This method employs a Gaussian-based modeling approach to approximate the point-spread function of each diffraction-limited object. This approach is used first to the position of objects in the primary channels (typically clathrin), followed by fitting Gaussian models at this position in the secondary channels (e.g. NCK1/2, PI3K, or EGF) to obtain the specific enrichment at each nanoscale clathrin object location. The amplitude of the Gaussian models is taken as the intensity of that channel’s protein within the detected object. To assess background and/or overlap occurring by random overlap of signals within the plasma membrane, the positions of secondary channels were randomized relative to the primary channel by rotating the image 180 degrees. Details on obtaining the software are available as described in (Lucarelli *et al*., 2017).

### Live cell time-lapse analysis of CCP dynamics

Automated detection, tracking and analysis of CCPs (as in **Figure 2**) was as previously described (Aguet *et al*., 2013; Delos Santos *et al*., 2017; Cabral-Dias *et al*., 2022; Rahmani *et al*., 2023). Image acquisition was performed, as described above under the heading *Fluorescence labeling and microscopy*, in ARPE19 cells stably expressing eGFP-clathrin light chain with cells maintained at 37C in phenol-red free DMEM/F12 media (Gibco) supplemented with 20 mM HEPES during imaging. Following time-lapse imaging at 1 Hz, diffraction-limited clathrin structures were detected using a Gaussian-based modeling approach (Aguet *et al*., 2013), and trajectories were determined from clathrin structure detections using u-track software (Jaqaman *et al*., 2008). sCLSs were distinguished from bona fide CCPs based on unbiased analysis of clathrin intensities in the early CCP stages (Aguet *et al*., 2013; Kadlecova *et al*., 2017). Both sCLSs and CCPs represent nucleation events, but only bona fide CCPs represent structures that undergo stabilization, maturation and in some cases scission to produce vesicles. We report the sCLS nucleation, CCP initiation, CCP lifetime distribution, and the density of persistent CCPs, as well as the intensity of eGFP-CLC within structures as the ‘plateau intensity’ of eGFP-clathrin within these structures.

### Cellular Proliferation and Viability Using Incucyte Imaging System

ARPE-19 cells were cultured in 6-well plates and subjected to siRNA transfection as specified. Following the second transfection, the cells were incubated in an Incucyte SX5 at 37°C with 5% CO2 (Sartorius, Göttingen, Germany) for continuous monitoring and automated quantification of cell growth and proliferation, as outlined in (Lo *et al*., 2022). Images were captured using a phase-contrast camera at 2-hour intervals over a period of 48 h. These time-lapse images were processed using the Incucyte Basic Analysis Software. For assessing cell confluence, phase contrast images were evaluated with a minimum area filter set to 25 μm² and a segmentation adjustment of 0.8. Confluence measurements were normalized to the baseline confluence observed in initial images. Statistical evaluations were conducted using two-way ANOVA and subsequent Tukey post-test, setting a significance threshold at p < 0.05 to identify differences between experimental conditions.

