## Supplemental Material for "NCK adaptor proteins regulate clathrin-coated pit dynamics and EGF-stimulated PI3K-Akt signaling"

This supplemental materials file contains the following:

Figure S1. Expression levels of Nck2-eGFP and Nck1-eGFP
Figure S2. Full images corresponding to images shown in Figure 1.
Figure S3. Knockdown validation of siRNA targeting Nck1 or Nck2.
Figure S4. Silencing of Nck1 and 2 with alternate siRNA targeting sequence in ARPE-19 cells.
Figure S5. Silencing of Nck1 and/or Nck2 in MDA-MB-231 cells decreases AKT activation and EGFR phosphorylation
Figure S6. Gab1 siRNA suppresses recruitment of PI3K p110β and EGFR to plasma membrane clathrin structures.

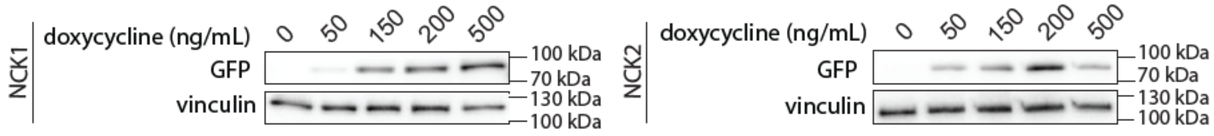


**Figure S1. Expression levels of Nck2-eGFP and Nck1-eGFP**. ARPE-19 cells carrying stable transgenes for inducible expression of Nck1-eGFP or Nck2-eGFP were treated with 0, 50, 150, 200, and 500ng/mL doxycycline for 24h. Shown are Western blotting of whole-cell lysates probed with anti-GFP at each doxycycline levels are shown with 0 doxycycline corresponding to no induction of either Nck1-eGFP or Nck2-eGFP.

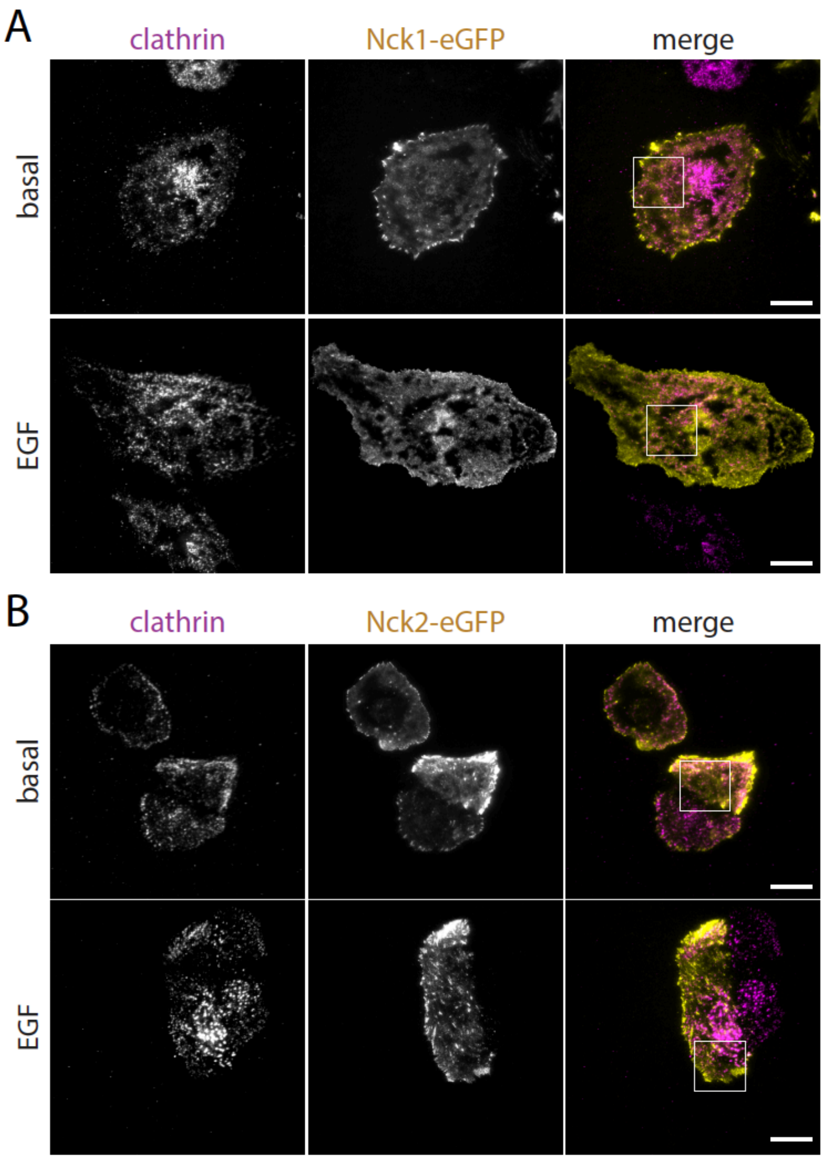


**Figure S2. Full images corresponding to images shown in Figure 1.** ARPE-19 cells carrying stable transgenes for inducible expression of Nck1-eGFP or Nck2-eGFP were treated with 150ng/mL doxycycline for 24h. Cells were stimulated with 20 ng/ml EGF for 5 min or left unstimulated (basal), then fixed and examined using TIRFM. Displayed (left) are representative images showcasing Nck1 or Nck2-associated clathrin structures. Scale = 20 µm. Boxes indicates image selections shown in **Figure 1**.

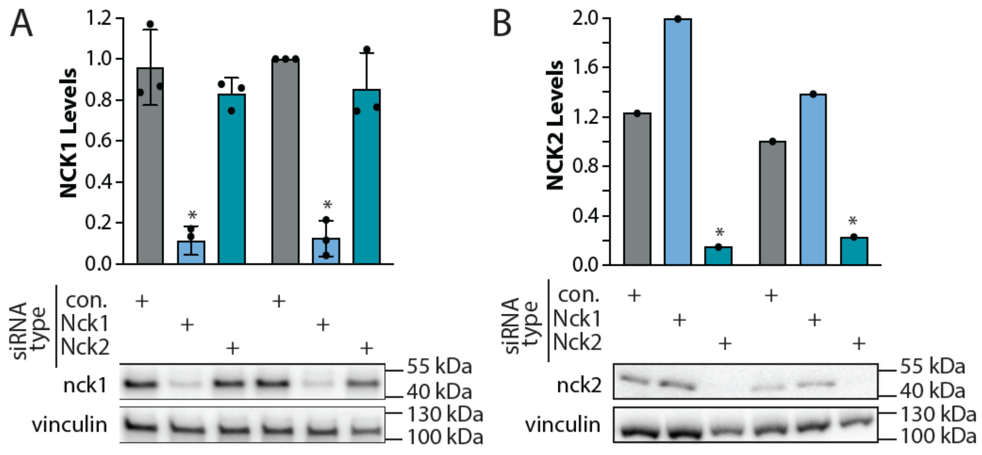


**Figure S3. Knockdown validation of siRNA targeting Nck1 or Nck2.**
ARPE-19 cells were transfected with siRNA targeting Nck1, Nck2, or nontargeting siRNA (control). **(A)** Western blotting of whole-cell lysates probed with anti-Nck1 or anti-Nck2 antibodies. **(B)** Shown are the mean ± SEM anti-Nck1 or anti-Nck2 with points representing individual experiment measurements; *, $P < 0.05$, relative to the control siRNA-treated conditions.

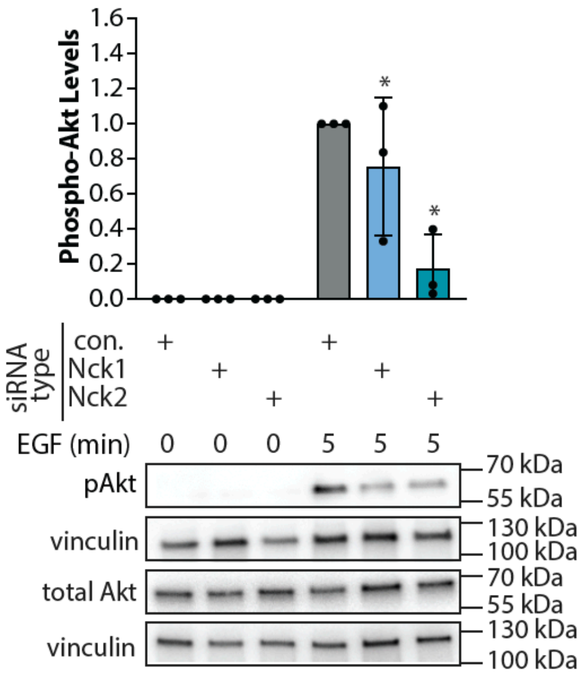


**Figure S4. Silencing of Nck1 and 2 with alternate siRNA targeting sequence in ARPE-19 cells.** ARPE-19 cells were transfected with siRNA targeting Nck1 (Nck1-2), Nck2 (Nck2-2), or nontargeting siRNA (control) using an alternate sequence to validate the specific siRNA targeting effect on Nck1 and Nck2, followed by stimulation with 5ng/ml EGF for 5 minutes. **(A)** Western blotting of whole-cell lysates probed with anti-phospho-Akt (Ser473). **(B)** Also shown are the mean ± SEM anti-phospho-Akt (Ser473) with points representing individual experiment measurements; n = 3; *, $P < 0.05$, relative to the control siRNA-treated EGF-stimulated condition.

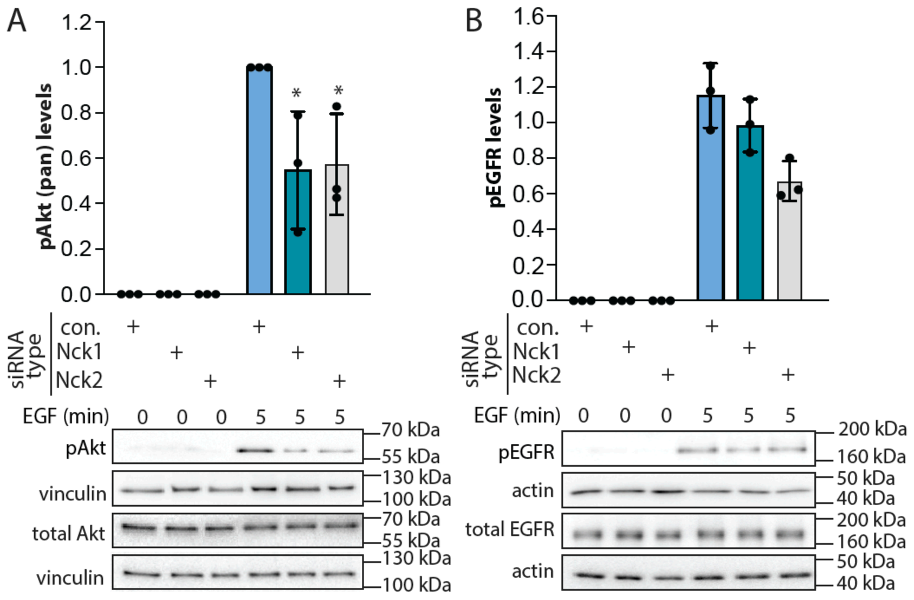


**Figure S5. Silencing of Nck1 or Nck2 in MDA-MB-231 cells decreases Akt activation.** MDA-MB-231 cells were transfected with siRNA targeting Nck1, Nck2, or nontargeting siRNA (control), followed by stimulation with 5ng/ml EGF for 5 min. **(A)** Western blotting of whole-cell lysates probed with anti-phospho-Akt (Ser473); also shown are the mean ± SEM phospho-Akt with points representing individual experiment measurements; *n = 3*; *, P < 0.05, relative to the control siRNA-treated EGF-stimulated condition. **(B)** Western blotting of whole-cell lysates probed using anti-phospho-EGFR (pY1068); also shown are the mean ± SEM phospho-EGFR with points representing individual experiment measurements; *n = 3*; *, P < 0.05, relative to the control siRNA-treated EGF-stimulated condition at 5 min.

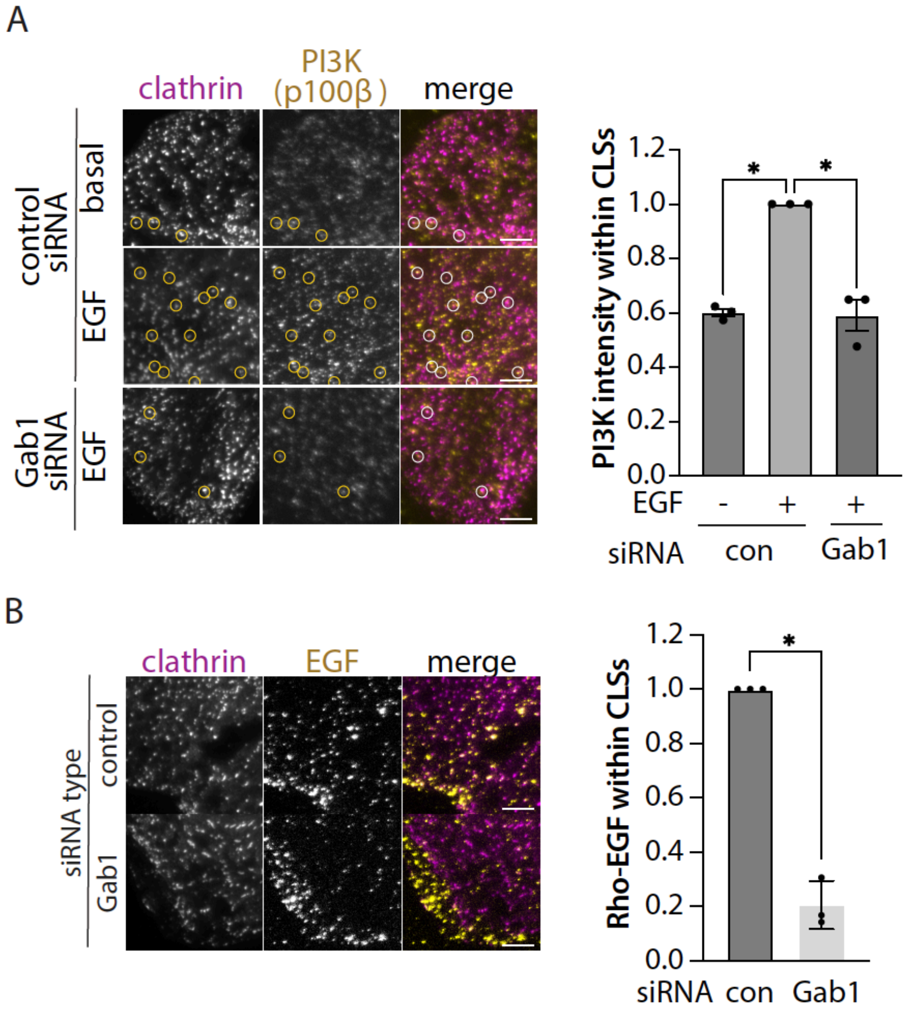


**Figure S6. Gab1 siRNA suppresses recruitment of PI3K p110β and EGFR to plasma membrane clathrin structures.** ARPE19 cells were subject to siRNA silencing as indicated. On the day of the experiment, cells were stimulated with 20 ng/mL EGF (***A***) or 20 ng/mL rhodamine-EGF (***B***) as indicated, followed by fixation; samples in (A) were also subjected to labeling of p110β (PI3K catalytic subunit) by antibody staining. Samples were then examined using TIRFM. Displayed (left) are representative images showing PI3K or rhodamine-EGF-associated clathrin structures. Scale = 5 μm. Images obtained by TIRFM underwent automated detection and analysis of CLSs, facilitating the quantification of PI3K or rhodamine-EGF enrichment within each identified structure. Shown are the fluorescence levels of rhodamine-EGF within CLSs from n = 3 independent experiments shown as mean ± SEM. The total number of CLSs and cells quantified (respectively) are as follows: (A) control siRNA basal: 9447, 55, control siRNA EGF-stimulated: 9986, 48; Nck1/2 siRNA EGF-stimulated: 7921, 61; (B) control siRNA: 17076, 60, Gab1 siRNA: 17420, 60; *, $p < 0.05$ determined by one-way ANOVA with Tukey post-hoc test.
